# The AraC/XylS protein MxiE and its co-regulator IpgC control a negative feedback loop in the transcription cascade that regulates type III secretion in *Shigella flexneri*

**DOI:** 10.1101/2022.04.20.488988

**Authors:** Joy A. McKenna, Monika MA. Karney, Daniel K. Chan, Natasha Weatherspoon-Griffin, Brianda Becerra Larios, Maria Carolina Pilonieta, George P. Munson, Helen J. Wing

## Abstract

Members of the <u>A</u>raC <u>F</u>amily of <u>T</u>ranscriptional <u>R</u>egulators (AFTRs) control the expression of many genes important to cellular processes, including virulence. In *Shigella* species, the type III secretion system (T3SS), a key determinant for host cell invasion, is regulated by the three-tiered VirF/VirB/MxiE transcription cascade. Both VirF and MxiE belong to the AFTRs and are characterized as positive transcriptional regulators. Here, we identify a novel regulatory activity for MxiE and its co-regulator IpgC, which manifests as a negative feedback loop in the VirF/VirB/MxiE transcription cascade. Our findings show that MxiE and IpgC down-regulate the *virB* promoter and hence VirB protein production, thus, decreasing VirB-dependent promoter activity at *ospD1*, one of the nearly 50 VirB-dependent genes. At the *virB* promoter, regions required for negative MxiE- and IpgC-dependent regulation were mapped and found to be coincident with regions required for positive VirF-dependent regulation. In tandem, negative MxiE- and IpgC-dependent regulation of the *virB* promoter only occurred in the presence of VirF suggesting that MxiE and IpgC can function to counter VirF activation of the *virB* promoter. Lastly, MxiE and IpgC do not down-regulate another VirF-activated promoter, *icsA*, demonstrating that this negative feedback loop targets the *virB* promoter. Our study provides insight into a mechanism that may reprogram *Shigella* virulence gene expression following type III secretion and provides the impetus to examine if MxiE and IpgC homologs in other important bacterial pathogens such as *Burkholderia pseudomallei* and *Salmonella enterica* serovars Typhimurium and Typhi coordinate similar negative feedback loops.

**IMPORTANCE:** The large AraC Family of Transcriptional Regulators (AFTRs) control virulence gene expression in many bacterial pathogens. In *Shigella* species, the AraC/XylS protein MxiE and its co-regulator IpgC positively regulate the expression of type III secretion system genes within the three-tiered VirF/VirB/MxiE transcription cascade. Our findings suggest a negative feedback loop in the VirF/VirB/MxiE cascade in which MxiE and IpgC counter VirF-dependent activation of the *virB* promoter, thus, making this the first characterization of negative MxiE- and IpgC-dependent regulation. Our study provides insight into a mechanism that likely reprograms *Shigella* virulence gene expression following type III secretion, which has implications for other important bacterial pathogens with functional homologs of MxiE and IpgC.

## INTRODUCTION

Many Gram-negative bacterial pathogens, including *Shigella* species, use type III secretion systems (T3SS) to directly inject virulence proteins, known as effectors, into host cells to invade and/or subvert host cell machinery [1]. Without a functional T3SS, these pathogens are often avirulent in animal infection models [2]. Frequently, genes encoding the T3SS are transcriptionally activated by members of the <u>A</u>raC <u>F</u>amily of <u>T</u>ranscriptional <u>R</u>egulators (AFTRs) [3, 4]. Examples of AFTRs that regulate T3SS-encoding genes are VirF [5, 6] and MxiE [7, 8] in *Shigella* species, BsaN in *Burkholderia pseudomallei* [9], and InvF in *Salmonella enterica* serovars Typhi and Typhimurium [10]. Of the well-characterized proteins in the large AraC family of transcriptional regulators (≥830 proteins; [11]), most are described as activators that regulate the transcription of genes involved in carbon metabolism, stress response, or pathogenesis [12]. A distinguishing feature of AFTRs is a conserved C-terminal DNA-binding domain comprised of two helix-turn-helix motifs [12, 13]. Due to the high homology of this domain, the DNA-binding and/or regulatory activities of some AFTRs have been shown to be interchangeable when substituted for each other [14–16]. In contrast, the N-terminal domains of AFTRs are functionally variable and involved in dimerization and/or ligand binding [4, 12, 17, 18]. Due to the notorious insolubility of purified AFTRs *in vitro*, functional characterizations for the mechanism(s) of AFTR transcriptional regulation have proven difficult [19].

In *Shigella flexneri*, a causal agent of bacillary dysentery, a T3SS is used to inject two distinct waves of effectors to invade human colonic epithelial cells and adopt intracellular residency [20–24]. Most genes encoding the T3SS (e.g., apparatus, effectors, and regulators) are clustered in the 31 kb *ipa mxi spa* operons [25, 26] on the large (~220 kb) virulence plasmid pINV [27–29]. The pINV-associated virulence genes are regulated by the three-tiered VirF/VirB/MxiE transcription cascade, of which the first- and third-tier regulators VirF and MxiE belong to the AFTRs. Upon a switch to 37 °C, *virF* expression is up-regulated [30–33], whereby, VirF transcriptionally activates *virB* [34–36]. VirB then counters transcriptional silencing mediated by the chromosomally encoded histone-like nucleoid structuring protein H-NS [37–45], which engages AT-rich DNA sequences [46–48], at pINV-associated genes. While 37 °C can be sufficient to relieve H-NS-mediated transcriptional silencing (e.g., *virF* [30–33, 49]), the expression of approximately 50 T3SS-encoding genes rely on VirB to counter silencing by H-NS [37, 39, 40, 42, 44]. The large VirB regulon [50] includes genes encoding the T3SS (e.g., secretion apparatus and first wave of effectors), other virulence associated factors (e.g., OspD1 [44], IcsP [40, 42, 43]), and the third-tier activator MxiE and its co-activator IpgC [51].

Prior to T3SS-dependent contact with the host cell, MxiE is sequestered by the anti-activator OspD1 and co-anti-activator/chaperone Spa15 [44, 52] whereas IpgC is independently sequestered by either the anti-co-activator IpaB or IpaC. Upon contact, the first wave of VirB-dependent effectors is secreted, which includes the anti-activator OspD1 and anti-co-activators IpaB and IpaC [7, 51, 53–55]. In doing so, MxiE becomes free to associate with the chaperone IpgC, which lacks a DNA-binding domain [56], and together MxiE and IpgC transcriptionally activate genes encoding the second wave of T3SS effectors [7, 8, 50, 56–59]. Due to the dormancy of MxiE- and IpgC-dependent transcriptional regulation prior to type III secretion, characterization of this complex regulatory system has proven challenging. In laboratory settings, active type III secretion can be induced to allow MxiE to associate with IpgC using chemicals (e.g., Congo Red [8], the bile salt sodium deoxycholate [60]) or a *S. flexneri* mutant that lacks the T3SS apparatus tip proteins IpaB or IpaD [53]. More recently, MxiE- and IpgC-dependent regulation has been more directly tested by tightly controlling the expression of *mxiE* and/or *ipgC* on plasmids in a *S. flexneri* strain cured of pINV [52]. Through a combination of these strategies, the expression of over a dozen pINV-associated (i.e., *ipaH7.8*, *ipaH9.8*, *ospB*, *ospC1*, *ospE1*, *ospF*, and *virA*; [8, 50, 58, 61]) and chromosomal (i.e., *ipaHa*, *ipaHc*, *ipaHd*; *gem1*, and *gem3* [58, 61]) genes have been demonstrated to be MxiE- and IpgC-dependent. By aligning these promoter regions, a 17 bp putative *cis*-acting MxiE regulatory site known as the MxiE Box was identified, which has subsequently been shown to be required for MxiE- and IpgC-dependent transcriptional activation [8, 58, 59]. However, as with many AFTRs, direct evidence for MxiE binding to a DNA recognition site remains elusive.

In this study, we identify and characterize a novel negative MxiE- and IpgC-dependent feedback loop in the three-tiered VirF/VirB/MxiE transcription cascade that regulates the expression of T3SS genes. We show that MxiE and IpgC negatively regulate the *virB* promoter, thus, decreasing VirB protein production and VirB-dependent promoter activity at *ospD1*. This is the first description of negative MxiE- and IpgC-dependent regulation. We propose and test a model for negative MxiE- and IpgC-dependent regulation, whereby, MxiE and IpgC interfere with VirF-dependent activation to negatively and specifically impact *virB* promoter activity. Our work has implications for other important bacterial pathogens with MxiE/IpgC homologs that control type III secretion systems, including BsaN/BicA in *Burkholderia pseudomallei* and InvF/SicA in *Salmonella enterica* serovars Typhi and Typhimurium [9, 10, 62].

## RESULTS

### MxiE- and IpgC-dependent regulation of the VirB-dependent *ospD1* promoter is negative and likely indirect

Previously [44], an investigation of the VirB-dependent *ospD1* promoter revealed a sequence similar to the MxiE Box in both composition, 13/17 nt match (bolded Fig. 1A) and position, 29 nt between this site and the upstream flank of the −10 promoter element of *ospD1* (Fig. 1A; [8, 58, 59]). This putative site also contained 8 out of the 9 strictly conserved nucleotides (underlined; Fig. 1A) demonstrated to be required for MxiE- and IpgC-dependent transcriptional regulation [58]. Since MxiE- and IpgC-dependent regulation of *ipaHa* only requires a site with a 14/17 nt match to the MxiE Box consensus, we reasoned that MxiE and IpgC might also positively regulate *ospD1*. To test this, the putative MxiE Box was mutated by site-directed mutagenesis in the context of our low-copy *lacZ* reporter for the *ospD1* promoter, pP*ospD1-lacZ* [44]. Activity of the *ospD1* promoter was measured in wild-type *S. flexneri* (2457T) and an isogenic *virB* mutant (AWY3) using β-galactosidase assays. To circumvent the low levels of free MxiE and IpgC [52, 63] associated with cells grown in broth where type III secretion is only weakly active (<5%) [53], L-arabinose inducible plasmid vectors carrying *mxiE* and *ipgC* (pBAD18-*mxiE-ipgC*) or no additional genes (pBAD18) were introduced into each cell background. These cells were then grown in the presence of 0.2% L-arabinose prior to promoter activity assays.

**Figure 1.**
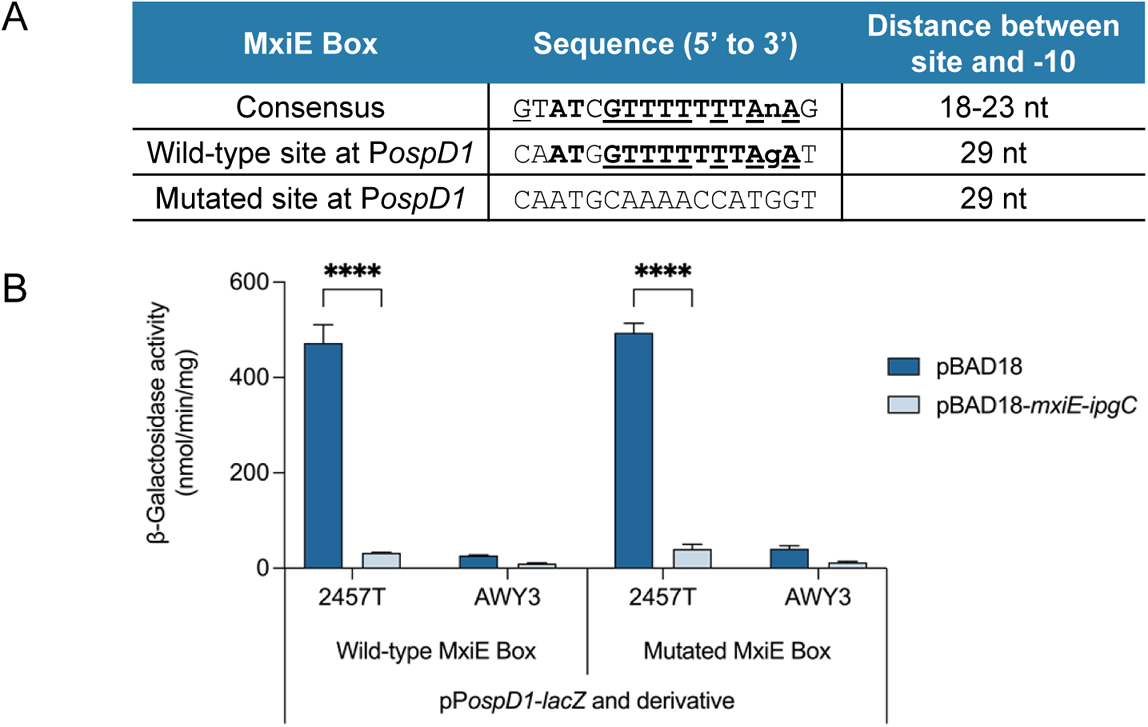
MxiE- and IpgC-dependent regulation of the VirB-dependent *ospD1* promoter is negative and indirect. **A)** Comparison of the sequence (in bold) and location of the MxiE Box consensus to the putative MxiE Box identified at the *ospD1* promoter (P*ospD1*). Nucleotides strictly conserved within the MxiE Box consensus [58] are underlined. Site directed mutagenesis was used to mutate the putative MxiE Box at P*ospD1*. **B)** Activities of the *ospD1* promoter were measured under inducing conditions (0.2% L-arabinose) in wild-type *S. flexneri* (2457T) and an isogenic *virB* mutant (AWY3) in the presence of exogenous MxiE and IpgC (pBAD18-*mxiE-ipgC*) or the empty control (pBAD18) using β-galactosidase assays. Representative data of three independent trials are shown. Data are represented as mean ± standard deviation. Significance calculated using two-way ANOVA with Šidák’s correction. * *p* < 0.0001.

Consistent with previous reports, *ospD1* promoter activity was VirB-dependent (18- to 19-fold change in +/− *virB* conditions; Fig. 1B; [44, 50]). However, while MxiE and IpgC were expected to increase *ospD1* promoter activity, this was not observed. Surprisingly, *ospD1* promoter activity decreased by 14- to 15-fold when *mxiE* and *ipgC* expression was induced compared to the pBAD18 empty control in wild-type *S. flexneri* (Fig. 1B). Furthermore, mutation of the putative MxiE Box did not impact *ospD1* promoter activity (Left vs. Right Panel; Fig. 1B) suggesting either indirect MxiE- and IpgC-dependent regulation of the *ospD1* promoter or the involvement of an unidentified MxiE Box. Since other putative MxiE Boxes with strong matches to the consensus were not identified in the *ospD1* promoter region contained within pP*ospD1-lacZ*, we concluded that the VirB-dependent *ospD1* promoter is negatively regulated by MxiE and IpgC and that this regulatory effect is likely to be indirect.

### The virB promoter is negatively regulated in a MxiE- and IpgC-dependent manner

From what is known regarding the VirF/VirB/MxiE transcription cascade, MxiE and IpgC are most likely to indirectly regulate the *ospD1* promoter by negatively impacting the transcription of *virB* (#1; Fig. 2) or *virF* (#2; Fig. 2). Prior transcriptomic analyses [50, 61] have shown that *virB* mRNA levels decrease both during constitutively active T3SS secretion (i.e., a condition favorable for free MxiE and IpgC; [52]) and in the presence of *mxiE* relative to the *mxiE* mutant. However, *virF* mRNA levels remained consistent in those cell backgrounds. As such, negative MxiE- and IpgC-dependent regulation most likely targets *virB* (#1; Fig. 2) rather than *virF* (#2; Fig. 2). Therefore, we hypothesized that *virB* is negatively regulated in a MxiE- and IpgC-dependent manner, thus, decreasing VirB protein production and VirB-dependent promoter activity like that of *ospD1*.

**Figure 2.**
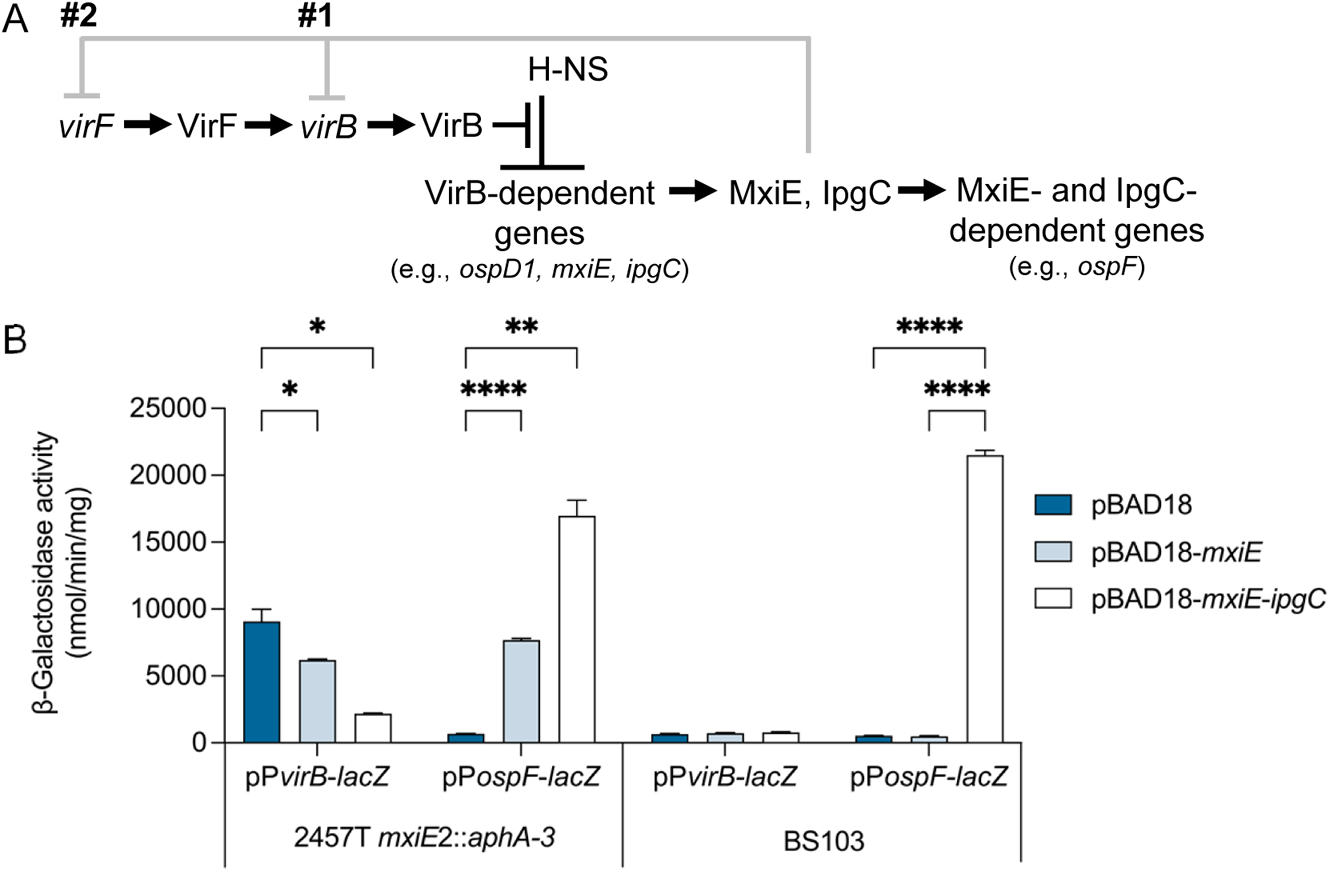
The *virB* promoter is negatively regulated in a MxiE- and IpgC-dependent manner. **A)** Potential transcriptional inputs for negative MxiE- and IpgC-dependent regulation at either *virB* (#1) or *virF* (#2) that may explain MxiE and IpgC down-regulation of the *ospD1* promoter. **B)** Differential MxiE- and IpgC-dependent regulation of the *virB* and *ospF* promoters. Promoter activities were measured using *lacZ* reporter plasmids, pP*virB*(−1946)-*lacZ* and pP*ospF-lacZ*, under inducing conditions (0.2% L-arabinose) in the *S. flexneri mxiE* mutant strain JAI04 (2457T *mxiE*2::*aphA-3*) and pINV-cured strain BS103 in the presence of exogenous MxiE (pBAD18-*mxiE*), MxiE and IpgC (pBAD18-*mxiE-ipgC*), or the empty control (pBAD18) using β-galactosidase assays. Representative data of three independent trials are shown. Data are represented as mean ± standard deviation. Significance calculated using two-way ANOVA with Tukey’s correction. * *p* < 0.05, ** *p* < 0.01, and **** *p* < 0.0001.

To address this hypothesis, we first determined if *virB* promoter activity is MxiE- and IpgC-dependent. To do this, the activity of the *virB* promoter was measured using β-galactosidase assays in a *S. flexneri mxiE* mutant strain JAI04 (2457T *mxiE2*::*aphA-3*) and the pINV-cured strain BS103 carrying either pBAD18-*mxiE*, pBAD18-*mxiE-ipgC*, or the pBAD18 empty control under inducing conditions. As a control for canonical MxiE- and IpgC-dependent transcriptional regulation, the *ospF* promoter was used since it has been well-established to be positively regulated by MxiE and IpgC (Fig. 2B; [8, 50, 58, 59]). As expected, *ospF* promoter activity significantly increased in the presence of exogenous MxiE or both MxiE and IpgC in the *mxiE* mutant cell background, exemplifying the previously described role of MxiE and IpgC as positive transcriptional regulators. In contrast, and yet under the same assay conditions, *virB* promoter activity significantly decreased when either *mxiE* or both *mxiE* and *ipgC* were induced compared to the empty pBAD18 control in the *mxiE* mutant cell background (Fig. 2B). Additionally, in the BS103 cell background that lacks pINV, where both *mxiE* and *ipgC* loci reside, we observed that *ospF* promoter activity only significantly increased when *mxiE* and *ipgC* were induced from pBAD18*-mxiE-ipgC*. Strikingly, however, basal *virB* promoter activity did not significantly change in the presence of *mxiE* or *mxiE* and *ipgC* (Fig. 2B). This suggests that negative MxiE- and IpgC-dependent regulation of the *virB* promoter requires an additional pINV-associated factor. Significantly, this is the first recorded observation of MxiE and IpgC either directly or indirectly negatively regulating transcription.

To address if MxiE and IpgC also decrease VirB protein levels, we measured VirB protein in wild-type *S. flexneri* (2457T), an isogenic *virB* mutant (AWY3), and an isogenic *mxiE* mutant strain JAI04 (2457T *mxiE2*::*aphA-3*) carrying either pBAD18 or pBAD-*mxiE-ipgC* using the same inducing conditions as our reporter assays. As expected, VirB protein was only detectable in the wild-type background but not the *virB* mutant background (15-fold difference in +/− VirB production based on average arbitrary densitometry units; Figs. 3A & B). Strikingly, in the presence of exogenous MxiE and IpgC, VirB protein was also undetectable (15-fold decrease in +/− MxiE and IpgC conditions in the *mxiE* mutant cell background; Figs. 3A & B). These data demonstrate that VirB protein levels drop precipitously in the presence of MxiE and IpgC, which is consistent with our observation that MxiE- and IpgC cause a decrease *virB* promoter activity. Additionally, since *mxiE* and *ipgC* mRNA expression is VirB-dependent [50], our findings suggest a negative feedback loop in the VirF/VirB/MxiE transcription cascade that regulates T3SS-encoding genes.

**Figure 3.**
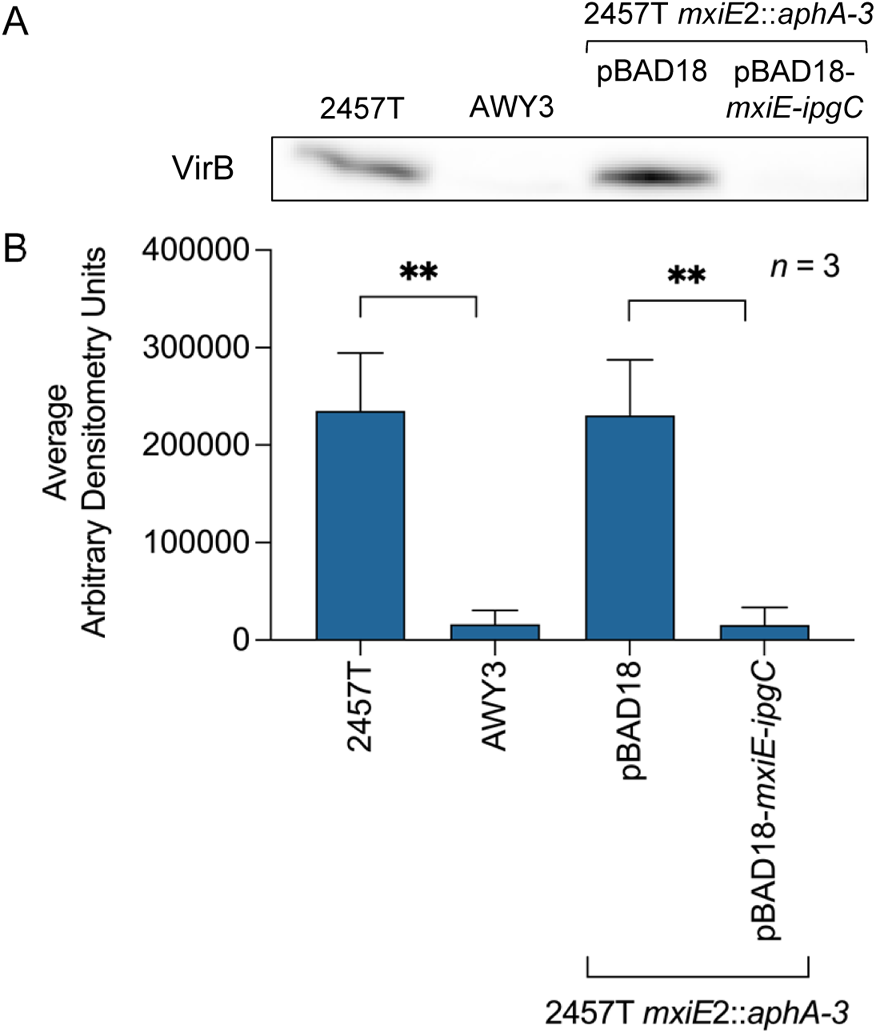
MxiE and IpgC decrease VirB protein levels. **A)** Western analysis of VirB in the presence (pBAD18-*mxiE-ipgC*) and absence (pBAD18) of induced MxiE and IpgC in the *S. flexneri mxiE* mutant strain JAI04 (2457T *mxiE*2::*aphA-3*). The *S. flexneri* wild-type (2457T) and isogenic *virB* mutant (AWY3) are positive and negative controls, respectively. Blot was probed using anti-VirB antibody (1:1000). A representative blot is shown. **B)** Densitometry analysis of Western analyses depicting the mean ± standard deviation of three independent trials (*n* = 3). Significance calculated using an unpaired two-tailed Student’s *t*-test that assumed either equal or unequal variance. ** *p* < 0.01.

### The regions required for negative MxiE- and IpgC-dependent regulation and positive VirF-dependent regulation of the *virB* promoter are coincident

To identify a potential mechanism for negative MxiE- and IpgC-dependent regulation at the *virB* promoter, the region upstream of *virB* was scanned for a putative MxiE Box [58, 59]. Four putative MxiE Boxes were identified (Site 1: 5’-AAATAGTAATTTTTaAG-3’, Site 2: 5’-GATAAGCATTTTTTcAT-3’, Site 3: 5’-CTGCCGATTCTCTTtCT-3’, Site 4: 5’-AGACTGATTTTTTAtCA-3’ centered at −52, −150, −299, and −873 relative to the +1 of *virB*, respectively). However, these putative MxiE Boxes were either located far upstream of the −10, exhibited low matches to the consensus sequence (7-10 nt match), or both. The lack of a traditional MxiE Box suggested that a different sequence and/or mechanism may be used for negative MxiE- and IpgC-dependent transcriptional regulation. As such, we mapped the region required for negative MxiE- and IpgC-dependent regulation of the *virB* promoter using 5’ promoter truncation analysis. The activities of the 5’ *virB* promoter truncations were measured using β-galactosidase assays in *S. flexneri mxiE* mutant strain JAI04 (2457T *mxiE2*::*aphA-3*) carrying pBAD18-*mxiE-ipgC* or the pBAD18 empty control under inducing conditions. The resulting data (Supplemental Fig. 1) were subsequently expressed as fold repression by MxiE and IpgC (Fig. 4B) and reveal that the down-regulation of the *virB* promoter by MxiE and IpgC is greatest (4- to 6-fold repression) when sequences upstream of −402 relative to the *virB* +1 are present. MxiE- and IpgC-dependent repression gradually drops from 3.5- to 2-fold with the removal of sequences between −402 and −116 relative to the *virB* +1 despite the overall increase in *virB* promoter activity (Supplemental Fig. 1). Strikingly, once the established region required for positive VirF-dependent regulation of the *virB* promoter is removed (−110 to −80 relative to the *virB* +1; Fig. 4A) the negative regulatory effect of MxiE and IpgC falls below 2-fold (Fig. 4B).

**Figure 4.**
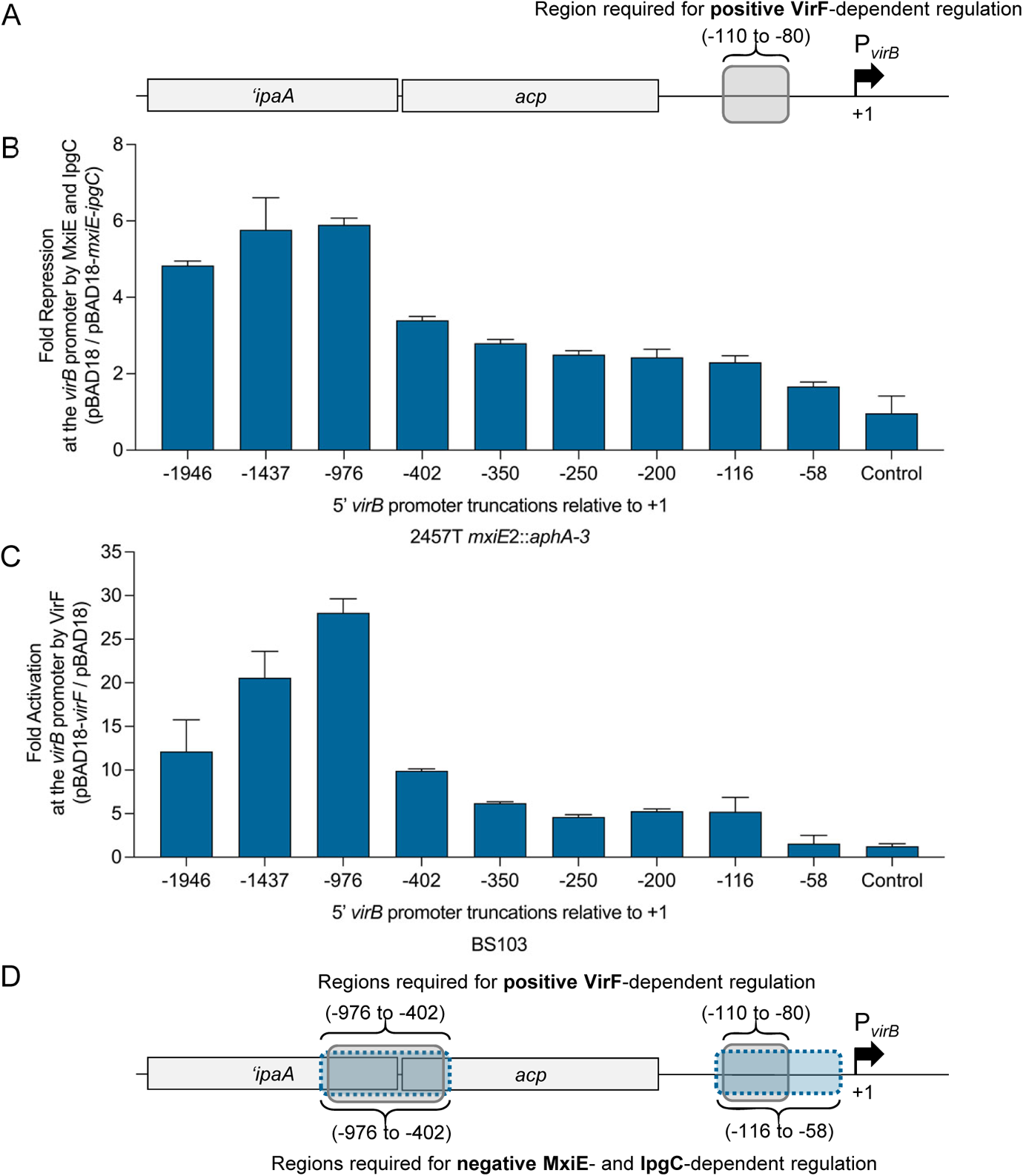
The regions required for negative MxiE- and IpgC-dependent regulation and positive VirF-dependent regulation of the *virB* promoter are coincident. **A)** Genetic locus of the *virB* promoter (−1946 to the *virB* +1). Drawn so panels align with corresponding 5’ *virB* promoter truncation in graphs below. **B)** Regions required for negative MxiE- and IpgC-dependent regulation of the *virB* promoter were mapped using 5’ promoter truncations. Activities of the 5’ *virB* promoter truncations were measured under inducing conditions (0.2% L-arabinose) in the *S. flexneri mxiE* mutant strain JAI04 (2457T *mxiE*2::*aphA-3*) in the presence of exogenous MxiE and IpgC (pBAD18-*mxiE-ipgC*) or the empty control (pBAD18) using β-galactosidase assays. Data are represented as the average in fold repression ± standard deviation (pBAD18/pBAD-*mxiE-ipgC*) in *virB* promoter activity from three independent trials. **C)** Regions required for positive VirF-dependent activation of the *virB* promoter were mapped using 5’ promoter truncations. Activities of the 5’ *virB* promoter truncations were measured under inducing conditions (0.2% L-arabinose) in the *S. flexneri* pINV-cured strain BS103 in the presence of exogenous VirF (pBAD18-*virF*) or the empty control (pBAD18) using β-galactosidase assays. Data are represented as the average in fold activation ± standard deviation (pBAD18-*virF*/pBAD18) in *virB* promoter activity from three independent trials. **D)** Schematic of the coincident regions required for negative MxiE- and IpgC-dependent regulation and positive VirF-dependent regulation of the *virB* promoter.

While the region required for VirF-dependent regulation of the *virB* promoter has been mapped [36] (Fig. 4A), those studies focused on a relatively short promoter region ~200 bp immediately upstream of the *virB* transcription start site (+1). Since our research has revealed long-range regulatory effects modulate genes encoded by the *Shigella* virulence plasmid, pINV [40, 42, 44], we re-examined VirF-dependent regulation of the *virB* promoter but with longer upstream promoter fragments. To do this, we measured VirF-dependent regulation of the *virB* promoter using the 5’ promoter truncations in the *S. flexneri* pINV-cured strain (BS103) carrying pBAD18-*virF* or the pBAD18 empty control under inducing conditions. The resulting data (Supplemental Fig. 2) were subsequently expressed as fold activation by VirF (Fig. 4C). As previously observed, VirF-dependent activation of the *virB* promoter is lost when DNA sequences upstream of −58 are removed, consistent with sequences between −110 to −80 being required for VirF-dependent regulation [36]. Surprisingly, VirF-dependent regulation was significantly higher when the upstream DNA sequences −1946 to −976 relative to the *virB* +1 were present. Even more striking was that the regions required for VirF-dependent regulation and MxiE- and IpgC-dependent regulation were coincident even though the regulatory effects were opposed (Figs. 4B-D; Supplemental Figs. 1 & 2). These data suggest that the AFTR MxiE and its co-regulator IpgC or a chromosomal MxiE and IpgC regulated factor interfere with the positive regulatory activity of VirF, another AFTR, from coincident regions upstream of the *virB* gene.

### Negative MxiE- and IpgC-dependent regulation of the *virB* promoter functions to counter VirF-dependent activation of *virB*

Since the regulatory regions for VirF and MxiE/IpgC at the *virB* promoter were superimposed, we next investigated if MxiE and IpgC negatively regulate *virB* by interfering with VirF-dependent activation of the *virB* promoter. To test this, MxiE- and IpgC-dependent regulation of the *virB* promoter was measured in a *S. flexneri* strain lacking pINV (BS103) but carrying either a *virF* (pBAD42-*virF*) or an empty expression plasmid (pBAD42). This strategy would allow these potentially complex and interconnected regulatory inputs to be assessed in a background without interference from other pINV-associated factors. As expected, *virB* promoter activity was positively regulated when VirF was present (pBAD18; Fig. 5A). In contrast, VirF-dependent *virB* promoter activity significantly decreased by 8- to 9-fold in the presence of MxiE and IpgC (pBAD-*mxiE-ipgC*) compared to the empty control (pBAD18; Fig. 5A). In the absence of *virF*, the activity of the *virB* promoter was not significantly altered regardless of the presence of *mxiE* and *ipgC* (Fig. 5A). These findings are consistent with data gathered in the pINV-cured BS103 cell background (Fig. 2B) since *virF* is encoded by pINV and consistent with the coincident pattern of VirF- and MxiE/IpgC-dependent regulation observed in the 5’ *virB* promoter truncation analysis (Fig. 4). Together, these data suggest that negative MxiE- and IpgC-dependent regulation of the *virB* promoter is mediated by these regulators interfering with VirF-dependent transcriptional activation of *virB*.

**Figure 5.**
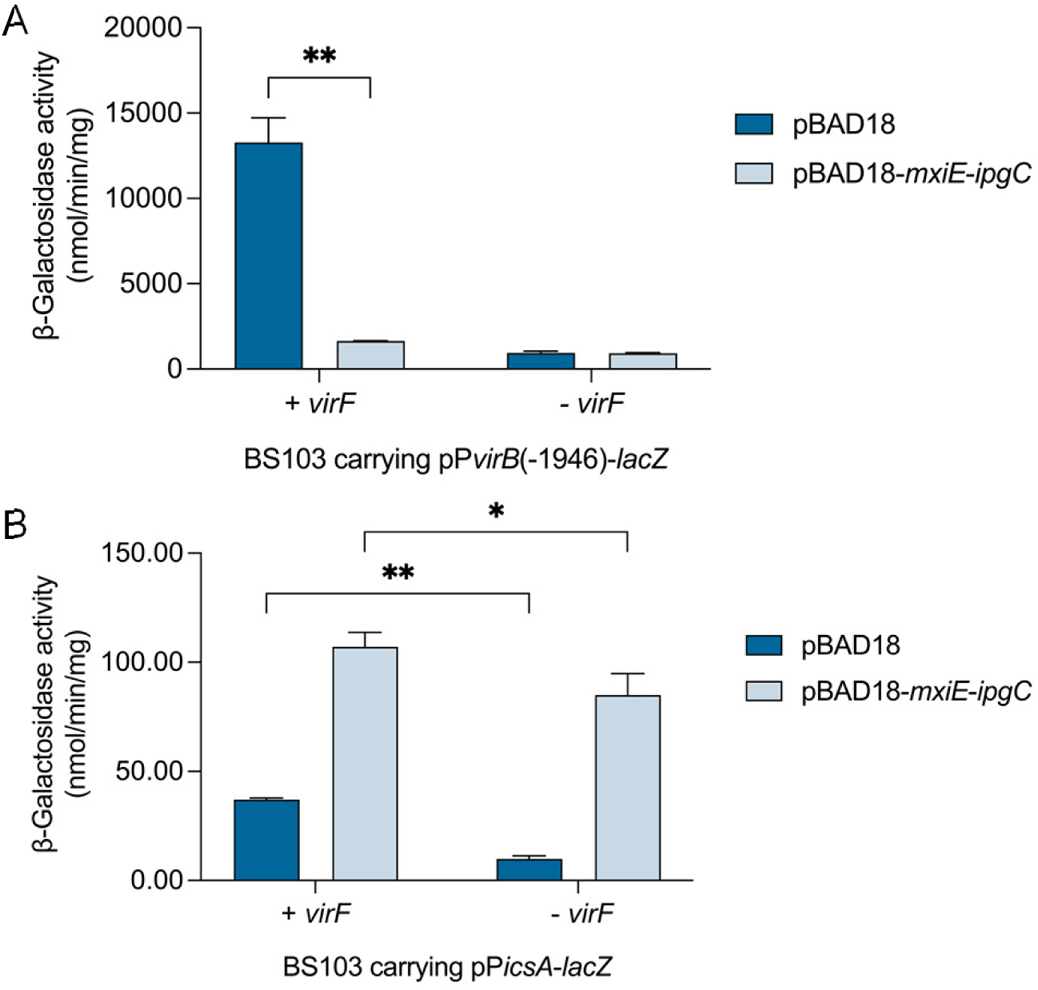
Negative MxiE- and IpgC-dependent regulation is observed at the VirF-dependent *virB* and not *icsA* promoter. **A)** Negative MxiE- and IpgC-dependent regulation of the *virB* promoter is solely observed in the presence of VirF. Activities of the *virB* promoter were measured from pP*virB*(−1946)-*lacZ* under inducing conditions (0.2% L-arabinose) in a *S. flexneri* strain cured of pINV (BS103) carrying either the pBAD42 empty vector or pBAD42-*virF* in the presence of exogenous MxiE and IpgC (pBAD18-*mxiE-ipgC*) or the empty control (pBAD18) using β-galactosidase assays. **B)** Activities of the *icsA* promoter were measured under inducing conditions (0.2% L-arabinose) in BS103 carrying either pBAD42 or pBAD42-*virF* in the presence of pBAD18-*mxiE-ipgC* or pBAD18 using β-galactosidase assays. Representative data of three independent trials are shown. Data are represented as mean ± standard deviation. Significance calculated using two-way ANOVA with Tukey’s correction. * *p* < 0.05 and ** *p* < 0.01.

### MxiE and IpgC do not negatively impact VirF-dependent activation of the *icsA promoter*

Lastly, we examined if MxiE and IpgC can counter VirF-dependent transcriptional activation of a different promoter. Since VirF has only been characterized to directly bind to and transcriptionally activate the pINV-associated *virB* [35, 36] and *icsA* [65–68] promoters, we tested if the *icsA* promoter was also negatively regulated by MxiE and IpgC. Activity of the well-characterized *icsA* promoter was measured in the presence (pBAD42-*virF*) and absence of *virF* (pBAD42) in a *S. flexneri* strain lacking pINV (BS103) that carries either pBAD18-*mxiE-ipgC* or the pBAD18 empty control under inducing conditions using β-galactosidase assays. Contrary to our results at the *virB* promoter, *icsA* promoter activity did not significantly decrease in the presence of MxiE and IpgC (pBAD-*mxiE-ipgC*) compared to the empty control (pBAD18) when in a BS103 cell background carrying pBAD42-*virF* (Fig. 5B). Rather, *icsA* promoter activity increased in a MxiE- and IpgC-dependent manner. Although the reason for this increase remains unclear, it may be caused by a MxiE Box within the *icsA-virA* intergenic region that is needed for positive MxiE- and IpgC-dependent regulation of *virA* [58, 59]. Regardless, these results demonstrate that MxiE and IpgC do not interfere with VirF-dependent activation of the *icsA* promoter, making it unlikely that MxiE and IpgC exert a dominant-negative effect on the VirF protein or lead to the expression of a negative regulator of VirF [70. Instead, the negative feedback loop characterized by this work and orchestrated by MxiE and IpgC, appears to down-regulate the activity of the *virB* promoter and hence the transcriptional cascade controlling *Shigella* virulence.

## DISCUSSION

Our study identifies a novel negative feedback loop in the VirF/VirB/MxiE transcription cascade that regulates the expression of T3SS genes in *S. flexneri.* We demonstrate that MxiE and IpgC, both VirB-dependent products, negatively regulate the *virB* promoter leading to decreased VirB protein levels and the down-regulation of *ospD1,* a representative of the large VirB regulon (>50 genes; [50]) (Fig. 6). This is the first description of negative regulation by MxiE and IpgC, which have been well established to function as transcriptional activators of genes encoding the second wave of T3SS effectors during *Shigella* infection [7, 8, 58, 59]. Additionally, our findings corroborate prior transcriptomics data that showed that both *ospD1* and *virB* expression increased in the absence of *mxiE* compared to wild-type [50].

**Figure 6.**
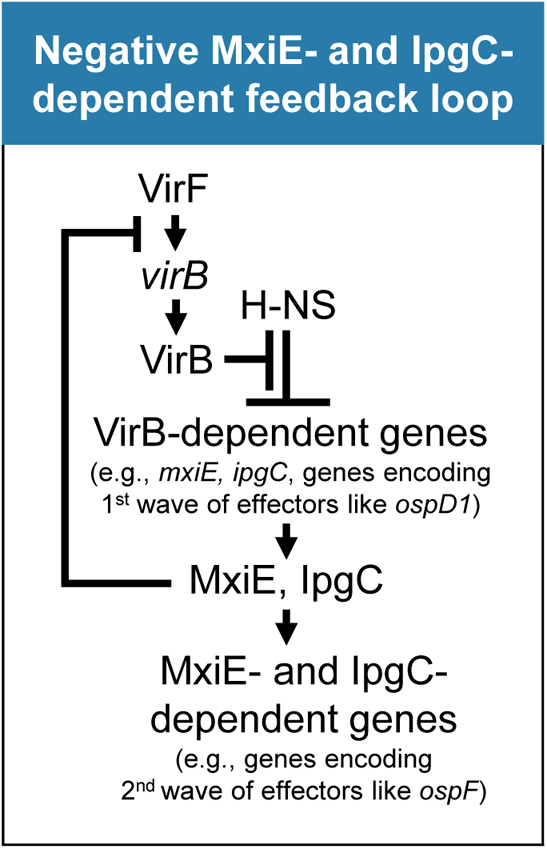
Model of negative MxiE- and IpgC-dependent feedback loop in the VirF/VirB/MxiE transcription cascade that regulates T3SS-encoding genes. The VirB-dependent third-tier regulators MxiE and IpgC negatively feedback at the *virB* promoter by countering VirF-dependent activation of the *virB* promoter. As a result, VirB-dependent promoter activity like that of *ospD1* is significantly decreased.

The negative feedback loop characterized in this work likely reprograms *Shigella* virulence gene expression at the stage following secretion of the first wave of T3SS effectors and invasion of host cells. This stage is favored since the secretion of the first wave of T3SS effectors triggers MxiE and IpgC to associate and positively regulate the transcription of second wave effector genes [7, 8, 52, 58, 59]. We report that when MxiE and IpgC levels rise in the cell, these proteins negatively regulate *virB* and in turn down-regulate VirB-dependent promoters such as *ospD1* (Fig. 1). Although not exhaustively tested in this study, the regulatory impacts of the MxiE- and IpgC-dependent negative feedback loop may be considerable. The VirB regulon consists of nearly 50 genes [50] including those that encode i) the T3SS secretion apparatus, ii) the first wave of effectors, iii) other virulence associated factors (e.g., OspD1 [44], IcsP [43]), and iv) the third-tier regulators MxiE and IpgC. Therefore, it is tantalizing to consider that negative MxiE- and IpgC-dependent regulation of *virB* may ‘tap the brakes’ on VirB-dependent gene expression, which could modulate or attenuate the expression of virulence genes in *Shigella* cells during an infection. Such an event could lead to an increase *in Shigella* persistence within host cells and/or the maintenance of *Shigella* virulence plasmid genes post-invasion (Note: prolonged VirB-dependent gene expression is energetically costly and can lead to pINV instability or loss [69]).

In addition, this work provides evidence for a potentially novel mechanism of negative AFTR-dependent regulation. We demonstrate that MxiE and IpgC negatively regulate the *virB* promoter in a manner dependent upon the transcriptional activator of the *virB* promoter VirF, which like MxiE also belongs to the AFTRs. Furthermore, we show that the regions required for negative MxiE- and IpgC-dependent regulation are coincident with the regions required for positive VirF-dependent regulation of the *virB* promoter. These data suggest that MxiE and IpgC may interfere with VirF-dependent activation, but specifically and only at the *virB* promoter. In support of this, our data show that the negative regulatory impact of MxiE and IpgC is solely observed at *virB* but not *icsA*, (the only other VirF-activated locus on pINV [65–68]). Thus, MxiE and IpgC are unlikely to interfere with VirF protein levels by exerting a dominant-negative effect on VirF or by up-regulating the production of an AraC Negative Regulator that targets VirF [70]. Instead, it is possible that MxiE and IpgC occlude VirF from the *virB* promoter to prevent VirF-dependent activation. Interestingly, a similar mechanism of AFTR regulatory interference has been described at the *rhaSR* operon in *E. coli* [71]. There, two AFTR members, RhaR and RhaS, recognize and bind to an overlapping region in the *rhaSR* promoter region. Interference can occur upon an increase in RhaS concentration that results in RhaS outcompeting RhaR for binding sites at the *rhaRS* promoter, thus, preventing the synergistic activation of this locus with another transcriptional activator, CRP [71]. While our work is consistent with a mechanism of AFTR interference being responsible for the MxiE- and IpgC-dependent negative feedback loop that controls the *Shigella* virulence gene cascade, *in vitro* experiments using all three purified proteins (MxiE, IpgC, and VirF) are required to gain direct evidence for this mechanism. Due to the notorious insolubility exhibited by AFTRs, this is currently beyond the scope of this study [7, 19, 57]. Nevertheless, our finding that the regions required for AFTR-mediated positive and negative regulation of the *virB* promoter are coincident is significant and may guide others when similar regulatory controls are detected.

While the specific sequence(s) responsible for negative MxiE- and IpgC-dependent regulation of the *virB* promoter was not identified in this work, our analysis suggested that this sequence(s) will not be organized like the MxiE Box consensus [58, 59]. Upon re-evaluation of the putative MxiE Boxes identified at the *virB* promoter region (Sites 1-4), it was found that Sites 1, 2, and 4 (−52, −150, and −873 relative to the +1 of *virB*, respectively) contained the direct repeat 5’-AnTTTTTnA-3’ which may constitute a half-site. Since AraC binds half-sites to loop DNA and negatively regulate transcription [72–74], it is possible that MxiE and IpgC negatively regulate via a similar mechanism. The significance of these sequences has yet to be determined but will frame future work that provides mechanistic insight into these opposing regulatory controls.

Finally, our finding that MxiE and IpgC create a negative feedback loop in the transcription cascade that regulates T3SS in *S. flexneri* has implications for MxiE and IpgC homologs that regulate T3SS-encoding genes in other bacterial pathogens like *Burkholderia pseudomallei* (i.e., BsaN and BicA [9]) and *Salmonella enterica* serovars Typhi and Typhimurium (i.e., InvF and SicA [10]). Of note, the MxiE homolog BsaN in *B. pseudomallei* has also been suggested to positively and negatively influence transcription in a transcriptomic analysis [75]. Therefore, other MxiE-like regulators, like BsaN or InvF, may also differentially regulate the expression of T3SS genes. We anticipate that other AFTR interference mechanisms may coordinate important negative feedback loops, like that described herein, to regulate virulence.

## EXPERIMENTAL PROCEDURES

### Bacterial strains, plasmids, and media

The bacterial strains and plasmids used in this study are listed in Table 1. *S. flexneri* strains were routinely grown at 25 or 37 °C in Luria-Bertani (LB) broth [76] with aeration by constant shaking (325 rpm in a LabLine/Barnstead 4000 MaxQ shaker) or on trypticase soy agar (TSA; trypticase soy broth containing 1.5% [w vol^-1^] agar). Where appropriate, antibiotics were used at the following final concentrations: ampicillin [100 μg ml^-1^], chloramphenicol [25 μg ml^-1^], kanamycin [50 μg ml^-1^], or spectinomycin [50 μg ml^-1^]. To ensure that *S. flexneri* strains maintained pINV during manipulation, Congo Red binding was tested prior to each assay on TSA plates containing 0.01% [w v^-1^] Congo Red (Sigma Chemical Co., St. Louis, MO.) at 37 °C.

**Table 1.**
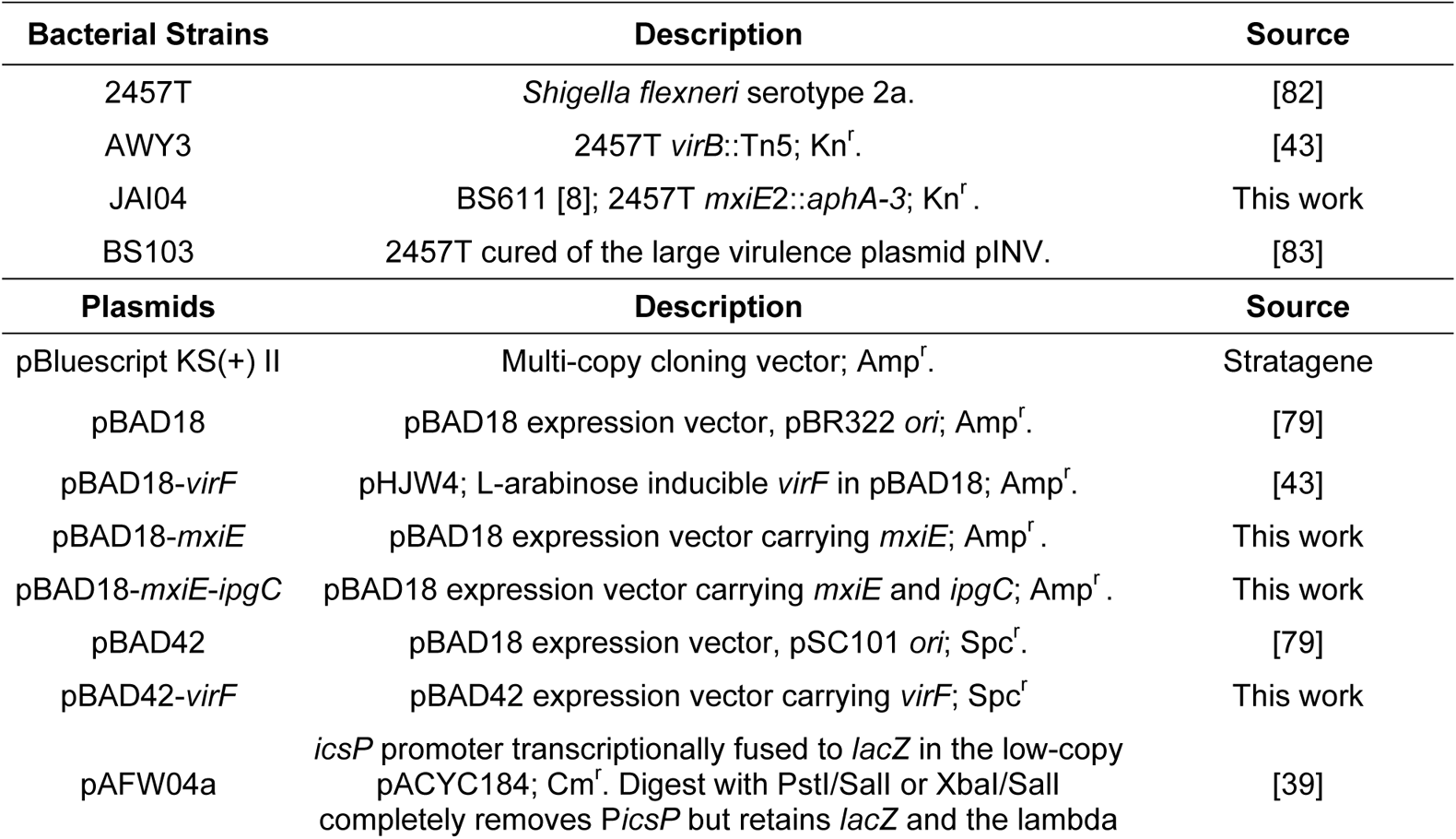
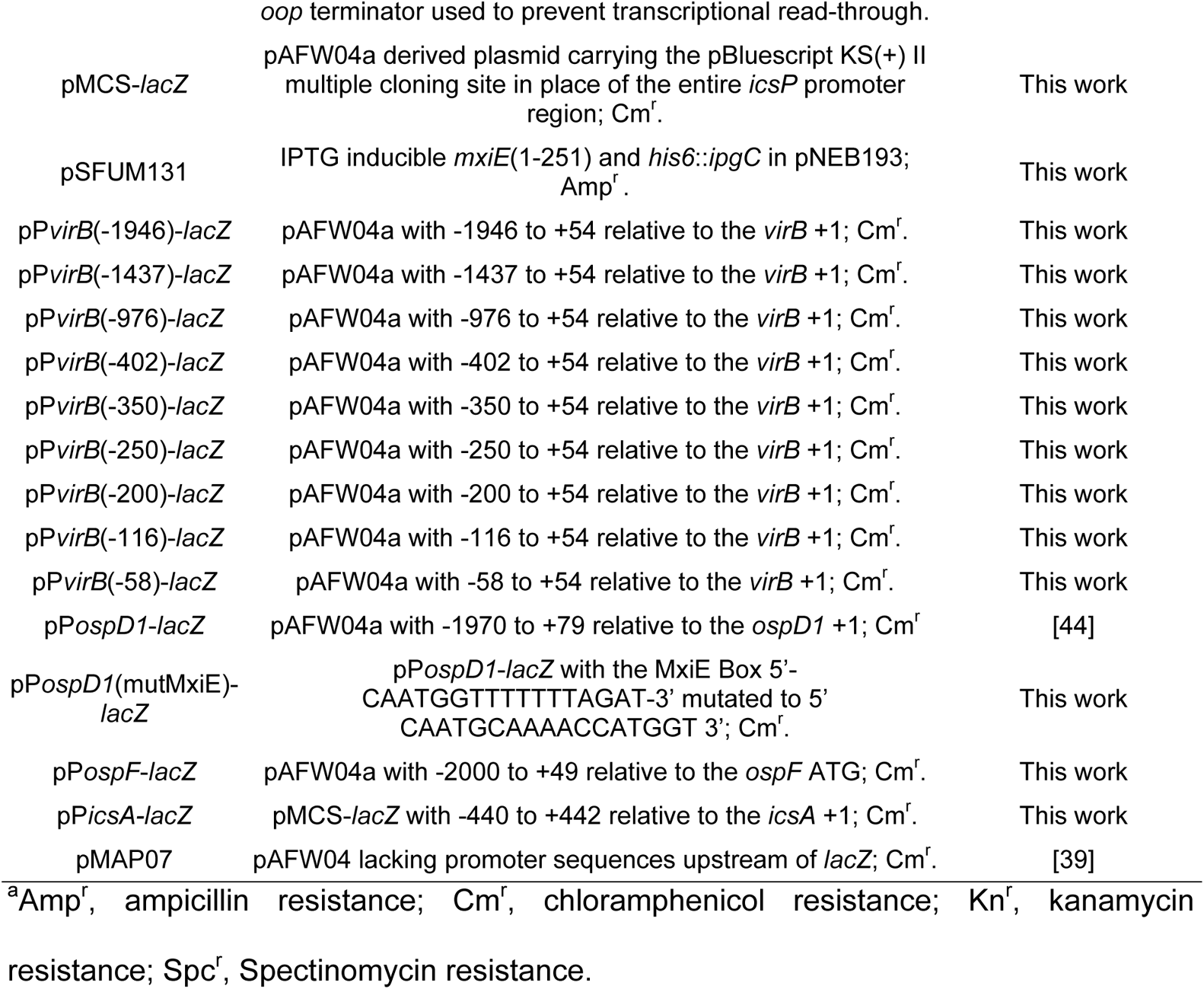
Bacterial strains and plasmids used in this study.

### Construction of *mxiE* mutant strain

To construct a 2457T *mxiE* mutant strain, *mxiE*2::*aphA-3* was transduced from BS611 [8] into 2457T by P1 phage transduction. The resulting strain, JAI04, was diagnosed by PCR for the *mxiE*2::*aphA-3* allele using primers W672 and W677. The integrity of pINV and key virulence loci [*icsA* (W11/W12), *virK* (W13/W14), *icsP* (W15/W16), *virF* (W17/W18), and *virB* (W19/W20)] were also verified by PCR. While BS611 is a 2457T derivative, JAI04 was created for lab strain consistency.

### Construction of the promoter-*lacZ* reporter plasmids

To create pP*ospD1*(mutMxiE)-*lacZ*, the putative MxiE Box 5’-CAATGGTTTTTTTAGAT-3’ was mutated to 5’-CAATGCAAAACCATGGT-3’ using site-directed mutagenesis in the context of pP*ospD1-lacZ* [44]. Briefly, pP*ospD1-lacZ* and gBlock 12 were digested with AflII and XbaI and then ligated to create pP*ospD1*(mutMxiE)-*lacZ*. An NcoI restriction site created within the mutated MxiE Box carries was exploited for the diagnostic verification.

To create the 5’ *virB* promoter truncations carrying −1946 to −58 relative to the *virB* +1, the upstream boundaries of *virB* were PCR amplified from the pINV of *S. flexneri* serotype 2a strain 2457T using the reverse primer W476 in combination with one of the following forward primers W696, W746, W747, W557, W558, W559, W560, W561, or W493 (Primers described in Table 2). Each PCR amplicon and the *lacZ* reporter plasmid, pAFW04 [39], were then digested with XbaI in combination with either PstI or SalI prior to ligation. All constructs were verified by DNA sequencing.

**Table 2.**
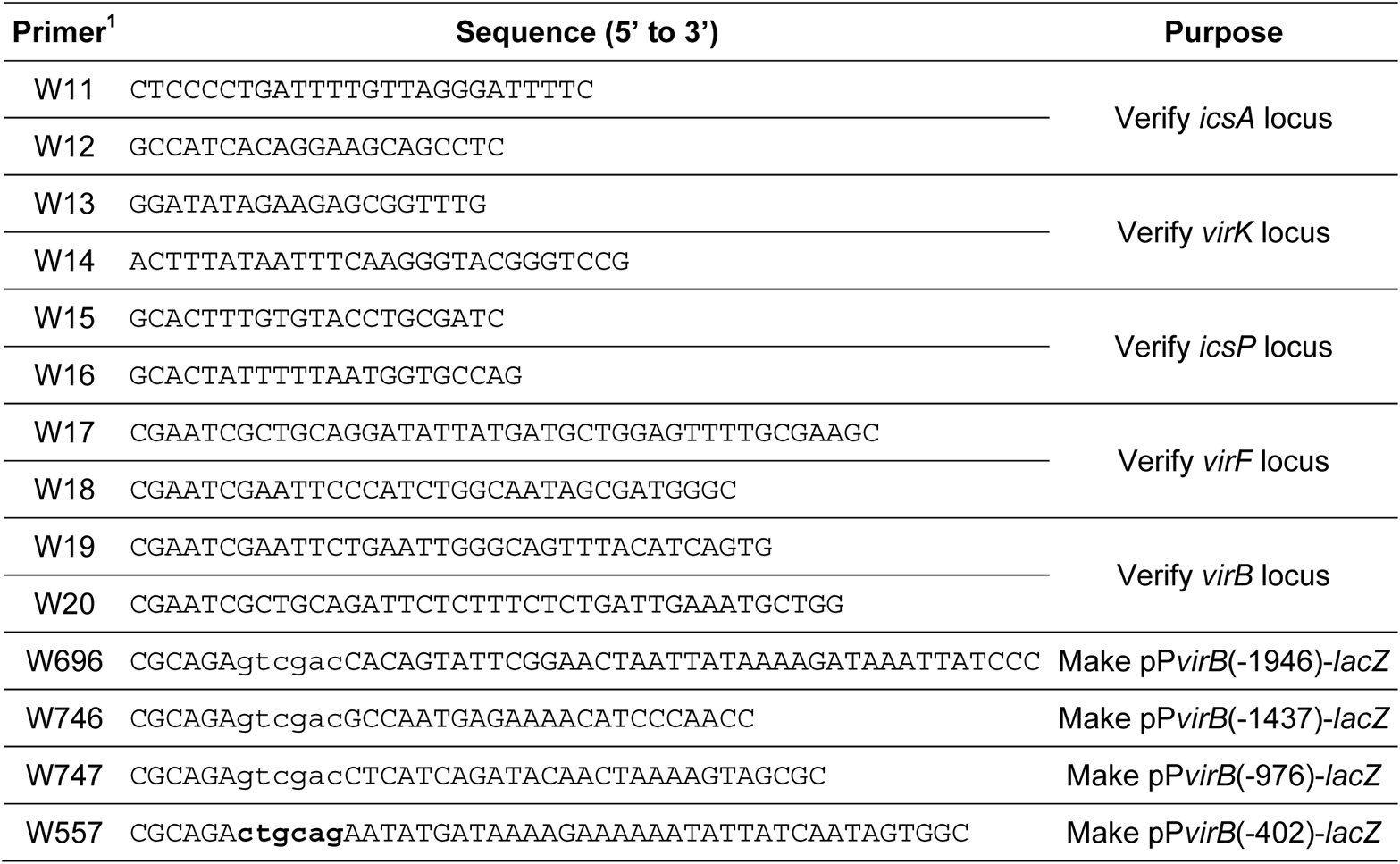
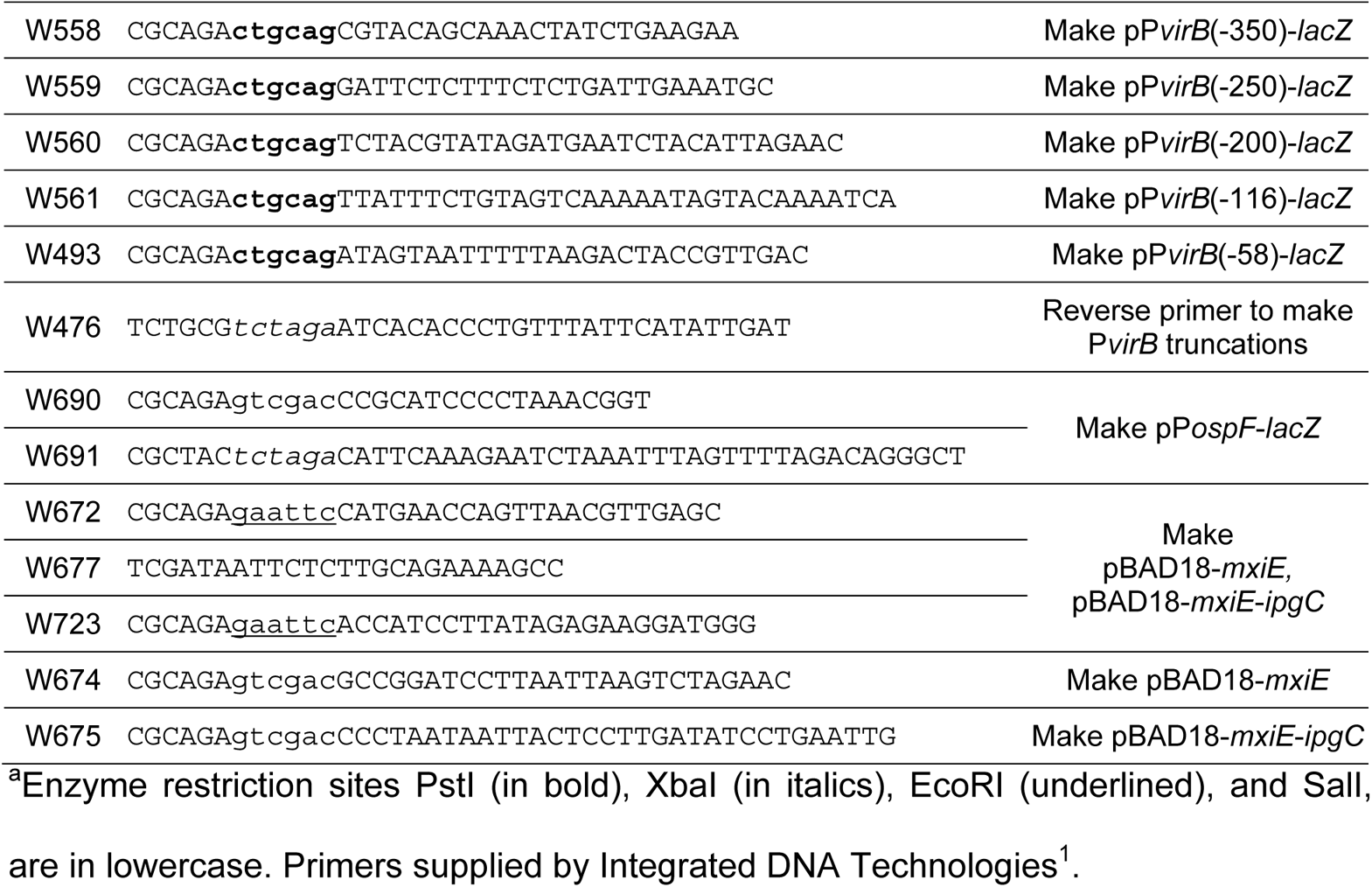
Oligonucleotide primers used in this study.

To create pP*ospF-lacZ*, the region spanning −1974 to +70 relative to the predicted +1 of *ospF* [59] was PCR amplified from the pINV of *S. flexneri* serotype 2a strain 2457T using W690 and W691. The PCR amplicon was then inserted into pBluescript KS(+) II by digesting both the amplicon and holding vector with XbaI and SalI prior to ligation. The amplified *ospF* regulatory region was verified by DNA sequencing. Subsequently, the *ospF* regulatory region was removed from pBluescript KS(+) II using XbaI/SalI and inserted into a similarly digested pAFW04 [39] to create pP*ospF-lacZ*, which was verified by diagnostic digest.

To create pMCS-*lacZ*, gBlock 14 (described in Table 3) carrying an engineered multiple cloning site was digested with PstI/XbaI. This insert was then ligated to pAFW04a [39], which had been digested with PstI and XbaI to entirely remove the *icsP* promoter region. The resulting construct, pMCS-*lacZ*, was verified by diagnostic digest.

**Table 3.**
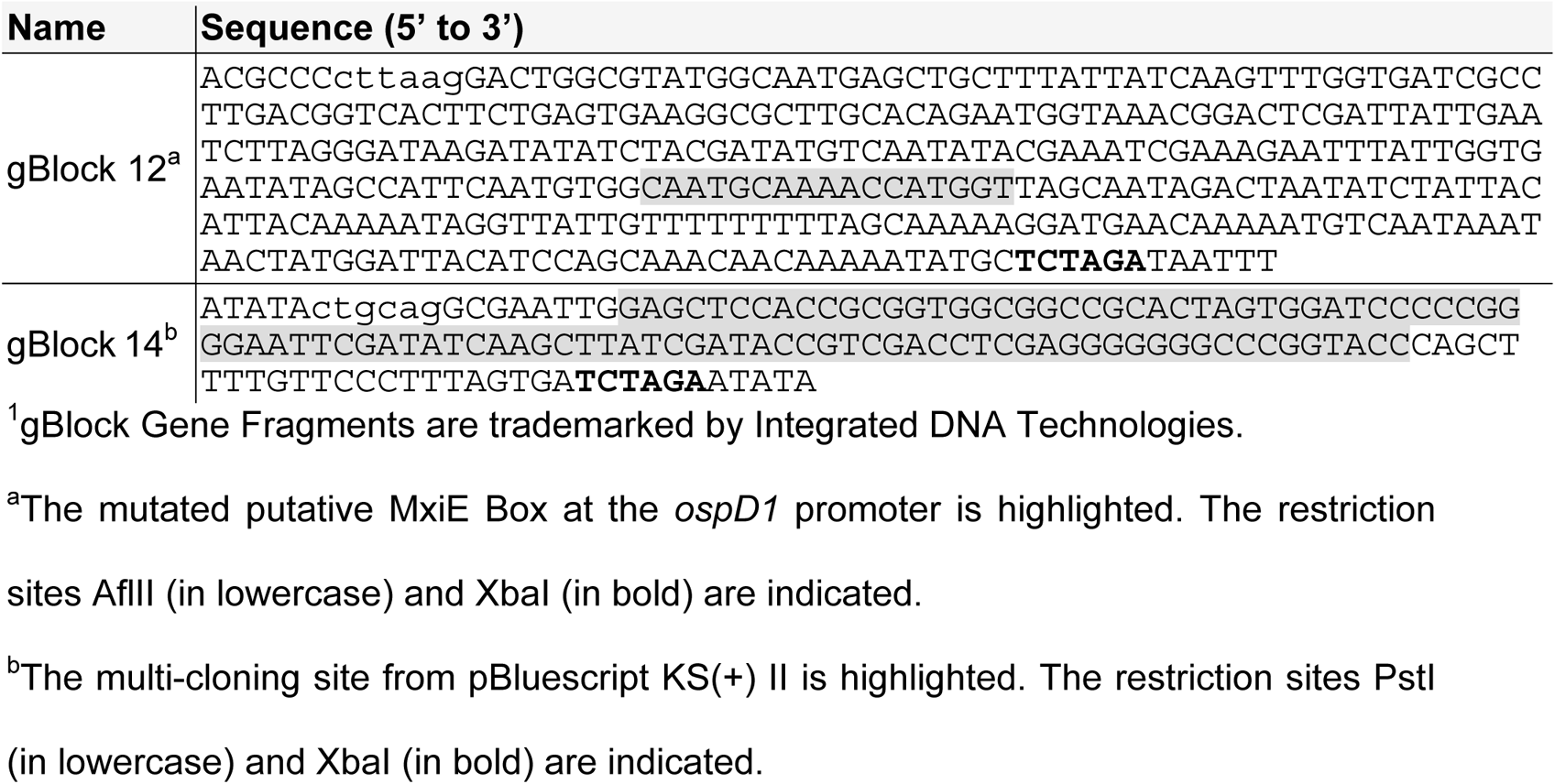
gBlock^1^ Gene Fragments used in this study.

To create pP*icsA-lacZ*, the region spanning −440 to +1610 relative to the +1 of *icsA* [66, 68] was PCR amplified from the pINV of *S. flexneri* 2a strain 2457T using W692 and W693. The PCR amplicon was then inserted into pBluescript KS(+) II by digesting both the amplicon and holding vector with XbaI and SalI prior to ligation. The entire inserted *icsA* regulatory region was verified by DNA sequencing. Then, the −440 to +442 region (includes all VirF and H-NS binding regions identified in [68]) relative to the *icsA* +1 was removed from pBluescript KS(+) II using XbaI and inserted into pMCS-*lacZ* linearized with XbaI to create pP*icsA-lacZ*, which was verified by diagnostic digest.

### Construction of protein expression plasmids

To create pBAD18-*mxiE* and pBAD18-*mxiE-ipgC*, *mxiE* and *ipgC* were PCR amplified from pSFUM131, which expresses the in frame and contiguous form of *mxiEab*(1-251) without transcriptional slippage and *his_6_*::*ipgC* from the IPTG inducible *lac* promoter in pNEB193 (Gift from Maria Carolina Pilonieta and George P. Munson; [77, 78]). Construction and verification of the pSFUM131 counterpart pSFUM139 (*malE*::*mxiE*(1-251) *his_6_*::*ipgC*) is described in [57]. To create pBAD18-*mxiE*, *mxiEab*(1-251) was amplified from pSFUM131 using W672 and W674. The PCR amplicon and pBAD18 [79] were then digested with EcoRI/SalI and ligated. The resulting construct was digested with EcoRI/BglII and ligated to a similarly digested PCR fragment amplified from pSFUM131 using W723 and W677. To create pBAD18-*mxiE-ipgC*, W672 and W675 were used to amplify *mxiEab* and *his_6_*::*ipgC* from pSFUM131. Then, the PCR amplicon and pBAD18 were digested with EcoRI/SalI and ligated. The resulting construct was digested with EcoRI/BglII and ligated to a similarly digested PCR fragment amplified from pSFUM131 using W723 and W677. These cloning strategies capture the alternative GTG translation start site required for *mxiEab*(1-251) [80].

To create pBAD42-*virF*, the *virF* gene was digested from pHJW4 [43] using EcoRI/SalI and ligated to a similarly digested pBAD42 [79]. The resulting construct was verified by digest with EcoRV.

All expression plasmids were verified by DNA sequencing and checked for regulatory activity and/or protein production by Western Blot.

### Quantification of promoter activity

Promoter activity was determined in a variety of strain backgrounds by measuring β-galactosidase activity (protocol adapted from [81] and described in [43]) from *lacZ* plasmid reporters (Table 1). Where indicated, *lacZ* reporter plasmids and pBAD18/42 derivatives were introduced into the different strain backgrounds (Table 1) by electroporation. All cultures were grown overnight (16 h) at 25 °C in LB broth [76] containing the respective antibiotics (Table 1) with aeration by constant shaking at 325 rpm. When using pBAD18, 0.2% D-glucose was added to the overnight LB broth. Prior to the β-galactosidase assay, overnight cultures were diluted 1:100 and grown for 5 h at 37 °C with constant shaking in LB broth with respective antibiotics prior to cell lysis. Where necessary, to induce expression from pBAD18 or pBAD42, 0.2% L-arabinose was added after 3 h of the 5 h growth period. Data represent β-galactosidase activities generated in triplicate.

### Protein analysis by Western Blot

Overnight cultures were diluted in 1:100 in LB containing the respective antibiotics and grown for 5 h at 37 °C with aeration by constant shaking at 325 rpm. To induce expression from pBAD18, 0.2% L-arabinose was added after 3 h of growth. Cells were normalized to cell density (OD_600_), washed with 0.2 M Tris buffer (pH 8.0), and resuspended in 10mM Tris (pH 7.4) containing 4x SDS-PAGE buffer and β-mercaptoethanol. VirB was detected using an affinity purified anti-VirB polyclonal antibody (1:1000; Pacific Immunology) and an anti-rabbit IgG-HRP (1:2000; GE; NA9340) secondary antibody. All blots were imaged on the Azure 500 (Azure Biosystems) and densitometry was completed using the AzureSpot analysis software.

### Statistical analysis

Statistical calculations were done using GraphPad Prism version 9.1.1 software. Statistical tests are indicated in the respective figure legends. Statistical significance is represented as * *p* <0.05, ** *p* <0.01, *** *p* <0.001, and **** *p* <0.0001.

## ACKNOWLEDGEMENTS

We graciously acknowledge Anthony T. Maurelli for gifting the *S. flexneri* strain BS611. We would also like to thank the following for their insightful discussions on this manuscript: Corrie S. Detweiler, Claude Parsot, Ronald K. Gary, and Michael A. Picker. This work was supported by the National Institute of Allergy and Infectious Diseases of the National Institutes of Health (NIH); R15 AI090573. DKC was supported by a grant from the National Institute of General Medical Sciences; GM 103440. We thank the UNLV Genomics Core Facility (sponsored by the National Institutes of General Medical Sciences; P20GM103440) for DNA sequencing services. This content is solely the responsibility of the authors and does not necessarily represent the official views of NIH. JAM has been a recipient of a Higher Education Graduate Research Opportunity Fellowship from the Nevada Space Grant Consortium NASA Training Grant NNX15AI02H and numerous fellowships and grants from UNLV and affiliated associations like the Association of Biology Graduate Students and the Graduate & Professional Student Association. These funders had no role in the study design, data collection and interpretation, or the decision to submit the work for publication. Lastly, we sincerely thank the anonymous reviewers for their comments and helpful suggestions.

## AUTHOR CONTRIBUTIONS

**Visualization, Writing-Original Draft Preparation**: JAM; **Conceptualization**: JAM, HJW; **Investigation and Validation**: JAM, MMAK; **Resources**: HJW, JAM, MMAK, DKC, NWG, BBL, MCP, GPM; **Supervision, Funding Acquisition**: HJW, JAM; **Writing – Review & Editing**: JAM, HJW, GPM, MMAK.

## FIGURE LEGENDS

**Supplemental Figure 1**. Regions required for negative MxiE- and IpgC-dependent regulation of the *virB* promoter were mapped using 5’ promoter truncations. Activities of the 5’ *virB* promoter truncations were measured under inducing conditions (0.2% L-arabinose) in the *S. flexneri mxiE* mutant strain JAI04 (2457T *mxiE*2::*aphA-3*) in the presence of exogenous MxiE and IpgC (pBAD18-*mxiE-ipgC*) or the empty control (pBAD18) using β-galactosidase assays. Representative data of three independent trials are shown. Data are represented as mean ± standard deviation. Each respective fold change (pBAD18/pBAD18-*mxiE-ipgC*) is shown.

**Supplemental Figure 2**. Regions required for positive VirF-dependent activation of the *virB* promoter were mapped using 5’ promoter truncations. Activities of the 5’ *virB* promoter truncations were measured under inducing conditions (0.2% L-arabinose) in the *S. flexneri* pINV-cured strain BS103 in the presence of exogenous VirF (pBAD18-*virF*) or the empty control (pBAD18) using β-galactosidase assays. Representative data of three independent trials are shown. Data are represented as mean ± standard deviation. Each respective fold change (pBAD18-*virF*/pBAD18) is shown.

